# Application of Multiomics Approach to Investigate the Therapeutic Potentials of Stem Cell-derived Extracellular Vesicle Subpopulations for Alzheimer’s Disease

**DOI:** 10.1101/2024.05.10.593647

**Authors:** Morteza Abyadeh, Alaattin Kaya

## Abstract

Alzheimer’s disease (AD) presents a complex interplay of molecular alterations, yet understanding its pathogenesis remains a challenge. In this study, we delved into the intricate landscape of proteome and transcriptome changes in AD brains compared to healthy controls, examining 788 brain samples revealing common alterations at both protein and mRNA levels. Moreover, our analysis revealed distinct protein-level changes in aberrant energy metabolism pathways in AD brains that were not evident at the mRNA level. This suggests that the changes in protein expression could provide a deeper molecular representation of AD pathogenesis. Subsequently, using a comparative proteomic approach, we explored the therapeutic potential of mesenchymal stem cell-derived extracellular vehicles (EVs), isolated through various methods, in mitigating AD-associated changes at the protein level. Our analysis revealed a particular EV-subtype that can be utilized for compensating dysregulated mitochondrial proteostasis in the AD brain. By using network biology approaches, we further revealed the potential regulators of key therapeutic proteins. Overall, our study illuminates the significance of proteome alterations in AD pathogenesis and identifies the therapeutic promise of a specific EV subpopulation with reduced pro-inflammatory protein cargo and enriched proteins to target mitochondrial proteostasis.

**Graphical Abstract:** 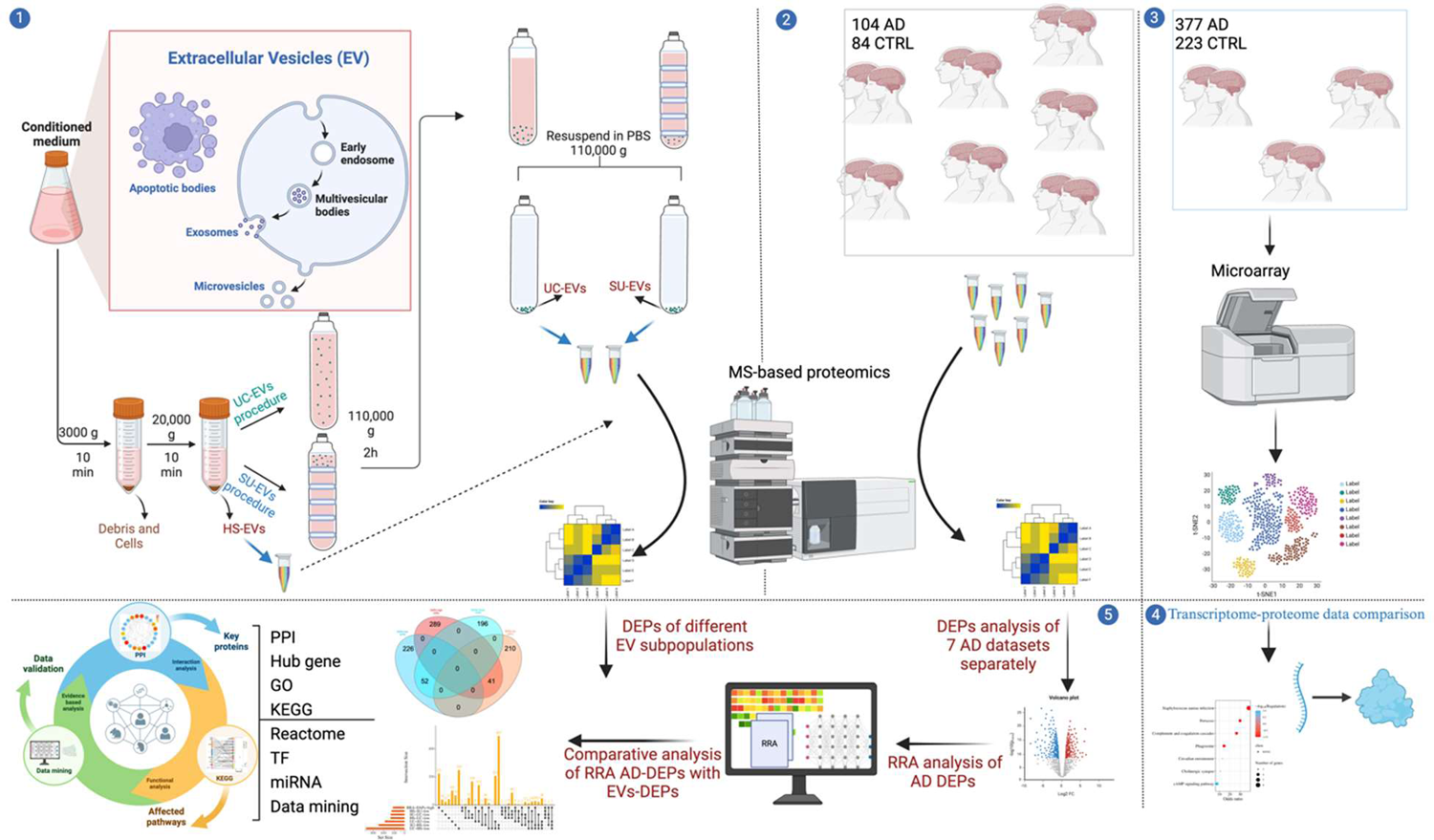

The cross-comparison of transcriptomics and proteomics data from AD brain samples and MSCs-derived EVs uncovers the therapeutic potential of a subpopulation targeting mitochondrial proteostasis and inflammation.

## Introduction

Alzheimer’s disease (AD) is a neurodegenerative, debilitating, and fatal disease characterized by the presence of extracellular plaques containing β-amyloid (Aβ) and intracellular neurofibrillary tangles (NFTs) containing tau proteins ^1^. AD is the most common type of dementia including nearly 70% of all cases ^2^. Age is the main risk factor for developing dementia and AD ^3^. Based on the World Health Organization report, currently 55 million people have dementia worldwide, and every year, there are nearly 10 million new cases ^4^. The total number of deaths attributable to AD has increased during the past 20 years, making it one the leading causes of mortality in the USA. According to the Alzheimer’s Association, almost 6.7 million Americans aged 65 and older are living with AD in 2023, which is projected to witness a dramatic increase, reaching an estimated 13 million by 2050 ^5^. The financial burden of AD and other dementia is estimated to be $345 billion in 2023 and almost $1 trillion in 2050 ^5^. This surge places a growing burden of emotional and financial challenges on Alzheimer’s patients, their loved ones, caregivers, and societies.

Despite significant advances in understanding the molecular pathogenesis of AD, a reliable treatment remains elusive. Recent optimism emerged in 2021 and 2023 with the introduction of AD antibodies (Aducanumab, Lecanemab) as a potential remedy ^6,7^. However, their efficacy is still under investigation, and their side effects and cost are notable. Consequently, there is a critical need for translational studies that can transform mechanistic insights into tangible impacts for AD patients and societies.

Mesenchymal stem cells (MSCs)-derived extracellular vehicles (EVs) have recently gained attention as promising therapeutic agents for various diseases, including neurodegenerative disorders. These EVs mimic the effects of stem cells while reducing the risk of risk of immune rejection and tumorigenicity, resulting from transplanted cells ^8^. EVs are small membranous structures released by almost all cell types that contain a diverse cargo of proteins, lipids, nucleic acids and metabolites. They can mediate intercellular communication and exert various biological effects, such as cell-to-cell signaling, immune modulation, and tissue repair ^9^. Three types of EVs, including microvesicles (MVs), exosomes, and apoptotic bodies are different in their biogenesis, release pathways, size, content, and function ^10^. EVs have been shown to reduce microglia activation, neuronal death, and improve blood-brain barrier integrity in AD animal models, that are all pathological signatures of AD ^8^.

There have been several lines of research on demonstrating exosomes as a reliable therapeutic option for various diseases, including neurodegenerative diseases ^8^. Particularly, therapeutic MSCs-derived exosomes have been shown to exhibit neuroprotective effects by regulating pathways associated with oxidative stress, neuroinflammation, apoptosis, reduced NFT and Amyloid plaques on cell and animal models of AD ^11,12^. Interestingly, large EVs have been shown to reduce pathological changes and restore homeostasis in a mouse model of AD more effectively ^13^. In addition, previous studies have highlighted that EVs isolated by different methods yield different size distribution and dependent on the size EVs carry distinct cargo that affects their features ^14,15^. These data suggests that isolation methods, and size should be considered in the selection of EV subpopulations based on their desired therapeutic application.

In this study, we conducted a meta-analysis using the robust rank aggregation method on publicly available proteomics (104 AD cases and 84 healthy controls) and transcriptomics (377 AD cases and 223 healthy controls) data from the frontal cortex of human brains. Our aim was to identify differentially abundant proteins (DAPs) and differentially expressed genes (DEGs) to perform a comparison between AD proteome and transcriptome data for examining whether protein changes are attributable to altered gene expression. As a result, we identified protein modules and disease-associated gene expression changes that were not directly observed at the mRNA level alone. The protein level robust changes were mostly associated with mitochondrial proteins indicating altered mitochondrial function in AD brains. In the second application, we performed a comparative analysis on proteomics data obtained from MSCs-derived different EV subpopulations with different size distribution isolated through different methods. Our analyses revealed particular EV subtypes that potentially be useful in restoring mitochondrial functions and alleviating inflammation in AD brains.

## Methods

### Proteome dataset selection and processing

A systematic search using “Proteome,” “Proteomic,” and “Alzheimer” was performed through the PubMed database. Studies included in AD brain proteome meta-analysis were selected if they met the following inclusion criteria: 1) reported protein expression in the frontal cortex region of AD patients and healthy controls; 2) both cases and controls were included. The protein expression data were extracted from published studies, and differentially abundant proteins (DAPs) were defined if the comparison of their expression in AD compared to controls was significant using one tail t-test (P-value < 0.05) without any fold change cutoff. Proteomic data from different subpopulations of EVs were retrieved from previously published papers. Daps for each EV subpopulation were identified using one-way ANOVA followed by a multiple comparison analysis with Tukey’s multiple comparison test in SPSS (26.0), and P-values <0.05 were considered statistically significant herein instead of doing multiple test correction analysis; we used log2 fold change cutoff > |0.263| as the additional criteria of defining DAPs ^16^. EVs were classified into three different subpopulations with different sizes and size distributions that were isolated through different isolation methods, including high-speed centrifuge (HS-EVs) with higher size distribution ranging from 50-1500 nm, ultracentrifuge (UC-EVs) with size distribution ranging from 100-300 nm and ultracentrifugation on sucrose cushion (SU-EVs) with size distribution of 300-1500 nm ^16^. The EV subpopulation was compared in binary mode, and we included two-sided comparisons to facilitate comparison with AD data. For example, the comparison between the HS vs SU group yielded the same differentially abundant proteins (DAPs) as the SU vs HS group but with reversed expression trends. DAPs that exhibited low abundance in the HS vs SU comparison were classified as high abundance in the SU vs HS comparison.

### Transcriptome dataset selection and processing

To analyze the transcriptome data from AD brains, the raw count files of brain transcriptome datasets related to AD versus healthy controls retrieved from the National Center for Biotechnology Information (NCBI) Gene Expression Omnibus (GEO) (https://www.ncbi.nlm.nih.gov/geo/), provided they satisfied the following criteria: 1) gene expression data reported for the frontal cortex of AD patients; and 2) inclusion of both AD cases and healthy controls. Differential expression analysis of genes (DEGs) between AD cases and healthy controls was conducted using the limma package. Genes with a p-value < 0.05 were considered significant and included for further analysis.

### Identification of robust DAPs and DEGs in AD and commonality with EV subpopulations

The updated “RobustRankAggreg” R package (Version: 1.2.1) was used to meta-analysis the AD brain proteome and transcriptome datasets and identify the robust DAPs and DEGs in AD brains^17^. To do this, the identified lists of low and high-abundant proteins from each dataset were separately ranked based on their fold changes. The list of low and high-abundant DAPs and down and upregulated DEGs from each separate dataset were combined into a single file and then subjected to the RRA method. Unlike the Venn diagram analysis, which identifies shared proteins, RRA identifies proteins with significant deregulation across datasets, even if they are not present in all ^18^. Identified robust DAPs and DEGs with a Bonferroni-corrected p-value less than 0.05 were considered statistically significant. Then, the list of robust DAPs was compared with the list of DAPs from different EV subpopulations using the “upset plot” to obtain common low- and high DAPs between AD and EV subpopulations. Moreover, the significance of the overlap between two protein datasets using both Fisher’s exact test and the hypergeometric test.

### Comparison of robust differentially abundant proteins with robust differentially expressed genes from AD brains

To assess whether molecular changes at the protein level are due to alteration in transcription or regulation beyond the gene expression, we compared the results of proteome analysis with transcriptome data. A comparison was performed between the list of down- and up-regulated robust DEGs and low- and high-abundant robust DAPs using Venn diagram analysis. Resulted common changes between the two datasets were then subjected to KEGG pathway analysis.

### Protein-protein interaction network analysis

Protein-protein interaction (PPI) networks were analyzed using the Cytoscape-String App plugin with a confidence score> 0.05, as previously described ^17^. Briefly, robust DAPs shared between AD and each EV subpopulation were uploaded into Cytoscape. Next, the Homo sapiens database in the StringDB was selected to reveal the protein interaction between differentially expressed proteins. Finally, to identify the hub genes within the protein network, CytoHubba, a plugin within Cytoscape, was utilized, and hub genes were selected based on the Maximal Clique Centrality (MCC) algorithm ^19^.

### Functional enrichment analysis

Gene ontology (GO) functional analyses, including Biological process (BP), Molecular function (MF), and Cellular component of common proteins between AD brain and EV subpopulations, were performed in Enrichr, a web-based tool for comprehensive gene set enrichment analysis (https://maayanlab.cloud/Enrichr/) ^20^. Biological pathway analysis was then performed in the KEGG (Kyoto Encyclopedia of Genes and Genomes) pathway and Reactome biological pathways database ^21,22^. In addition, ShinyGo, a web-based tool for comprehensive gene set enrichment analysis, was utilized to obtain the enrichment folds (http://bioinformatics.sdstate.edu/go/, ShinyGO 0.77) ^23^. Enriched terms with an adjusted p-value less than 0.05 were considered statistically significant for low-high abundant DAPs.

### Gene–miRNA and gene-transcription factors co-regulatory analysis

The NetworkAnalyst online tool was used to retrieve the hub gene—miRNA interactions network, which uses collected data of validated miRNA-gene interaction from TarBase and miRTarBase ^24^. In addition, the hub gene—miRNA and transcription factor (TF) coregulatory network analysis was retrieved from the NetworkAnalyst tool and then visualized using Cytoscape.

### Literature search for the validation of the identified miRNAs and TFs via text mining

To search miRNAs and TFs from our analysis and their relationship with AD, based on the published literature, the “batch_pubmed_dowload” function from the easyPubMed R package was employed. The articles, including our identified targets, were thoroughly discussed in the discussion. Resulted data from all steps of the analysis were visualized using Cytoscape (version 3.10.2), R (Version 2023.09.1+494), and Python (version 3.12.0).

## Results

### AD brain proteome meta-analysis revealed the shared key protein level changes

In total, seven proteome datasets including 104 AD and 84 healthy controls samples were included in our analysis (**Table 1**) ^25–28^. DAPs analysis showed significant differences between datasets in the terms of number of DAPs (**Figure 1A**). In this regard, individual case TMT analysis of Mount Sinai cohort with 39 AD cases and 23 controls showed higher number of total DAPs including 1279 low and 2114 high abundant proteins, and data from Goizueta Alzheimer’s Disease Research Center (ADRC) including 10 AD and 10 controls with 973 low and 564 high abundant proteins, showed lowest number of total DAPs (**Figure 1A**, **Supplementary file 1**). Interestingly, the total number of differentially abundant proteins (DAPs) does not necessarily reflect the total count of identified proteins or the number of included AD cases and controls. For instance, in the Pooled TMT analysis of the Banner Sun cohort with 36 participants, the highest number of identified proteins was observed, totaling 14,513, with 2,079 identified as DAPs. Conversely, the Goizueta Alzheimer’s Disease Research Center (ADRC), which included 19 participants, identified a total of 11,244 proteins, with 2,885 identified as DAPs (**Supplementary file 1**). In addition, number of differentially high abundant proteins were higher than those with low abundant in almost all studies. Accordingly, robust rank aggregation analysis also yielded the same distribution across robust DAPs including 277 low and 330 high abundant DAPs (**Figure 1B**, **Supplementary file 1**). Functional enrichment analysis of robust DAPs revealed the oxidative phosphorylation pathway, retrograde endocannabinoid signaling (which shares most of the proteins with oxidative phosphorylation pathway) and glutamatergic synapse among top downregulated pathways and focal adhesion, complement and coagulation cascades, and glycine, serine and threonine metabolism among top upregulated pathways (**Figure 1C**, **Supplementary file 2**). Of these enriched significant KEGG pathways, oxidative phosphorylation showed the highest odds ratio (OR: 32.85), indicating the key role of mitochondrial dysfunction in AD across different datasets (**Figure 1C**).

**Figure 1.**
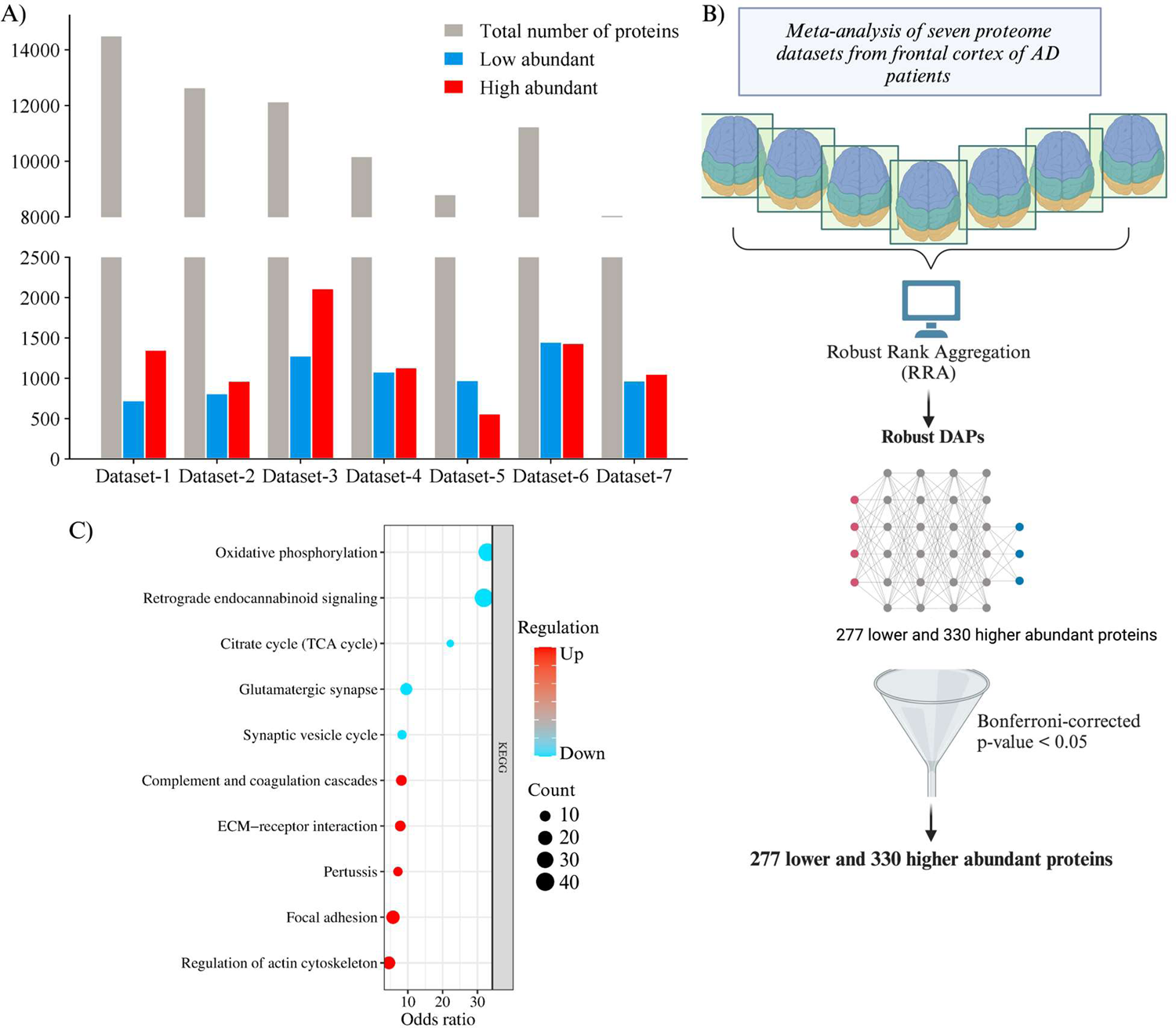
Proteomics features of AD datasets. A). Number of total, low, and high abundant DAPs in each dataset. B) Schematic summary of meta-analysis using RRA method performed on proteomic datasets; C). Top KEGG pathways enriched by robust DAPs in AD brains. DAPs, differentially abundant proteins; AD, Alzheimer’s disease; RRA, Robust rank aggregation.

**Table 1.** Characteristics of the selected proteomics datasets analyzed in this study.

| Number | Publication | Sources | Number of Proteins | Number of AD/CTRL | Age (years) AD/CTRL | Postmortem Interval (h) AD/CTRL |
| --- | --- | --- | --- | --- | --- | --- |
| Set-1 | Bai et al. Neuron 2020<sup>25</sup> | Pooled TMT analysis of Banner Sun cohort | 14513 | 18/18 | 75.7±8.7/83.3±8.0 | 3.3±1.0/2.8±0.6 |
| Set-2 | Bai et al. Neuron 2020<sup>25</sup> | Individual case TMT analysis of Banner Sun cohort | 12650 | 15/12 | 78.0±8.6/82.9±7.4 | 3.1±0.9/2.6±0.5 |
| Set-3 | Bai et al. Neuron 2020<sup>25</sup> | Individual case TMT analysis of Mount Sinai cohort | 12147 | 39/23 | 84±10.1/81±10.4 | 6.4±4.4/9.5±6.3 |
| Set-4 | Wang et al. Anal. Chem. 2020<sup>26</sup> | 16-plex TMT analysis of Banner Sun cohort | 10175 | 8/7 | 57±5.7/57.7 ± 5.2 | 3.0±0.8/NA |
| Set-5 | Higginbotham et al. Sci. Adv. 2020<sup>27</sup> | Goizueta Alzheimer's Disease Research Center (ADRC) | 8817 | 10/10 | 65.1±7.4/63.7±10.3 | 5.4±1.5/9.3±7.1 |
| Set-6 | Higginbotham et al. Sci. Adv. 2020<sup>27</sup> | Goizueta Alzheimer's Disease Research Center (ADRC) | 11244 | 9/10 | 65.2±7.9/63.7±10.3 | 5.5±1.6/9.3±7.1 |
| Set-7 | Sathe et al. J. Neurochem. 2021<sup>28</sup> | Baltimore Longitudinal Study of Aging (BLSA) cohort | 8066 | 5/4 | 87 ± 5.7/88 ± 6.8 | NA |

### Proteome and transcriptome comparison revealed molecular changes at protein level that was not identified at gene expression level

Next, to investigate the molecular changes in AD at both gene expression and protein translation levels we analyzed differentially expressed genes (DEGs) in AD brains and analyzed correlation between the robust DAPs and DEGs. We obtained three transcriptome datasets (GSE118553 ^29^, GSE48350 ^30^, and GSE33000^31^), in total 377 AD patients and 223 healthy controls brain samples were included in our analysis (**Table 2**). RRA analysis of transcriptome data returned 248 down and 252 up regulated significant DEGs. The comparison between robust DEGs and DAPs yielded 52 genes that are down regulated at both transcript and protein level. On the other hand, 41 genes were found to have high transcript and protein abundance in AD brains (**Figure 2A**).

**Figure 2.**
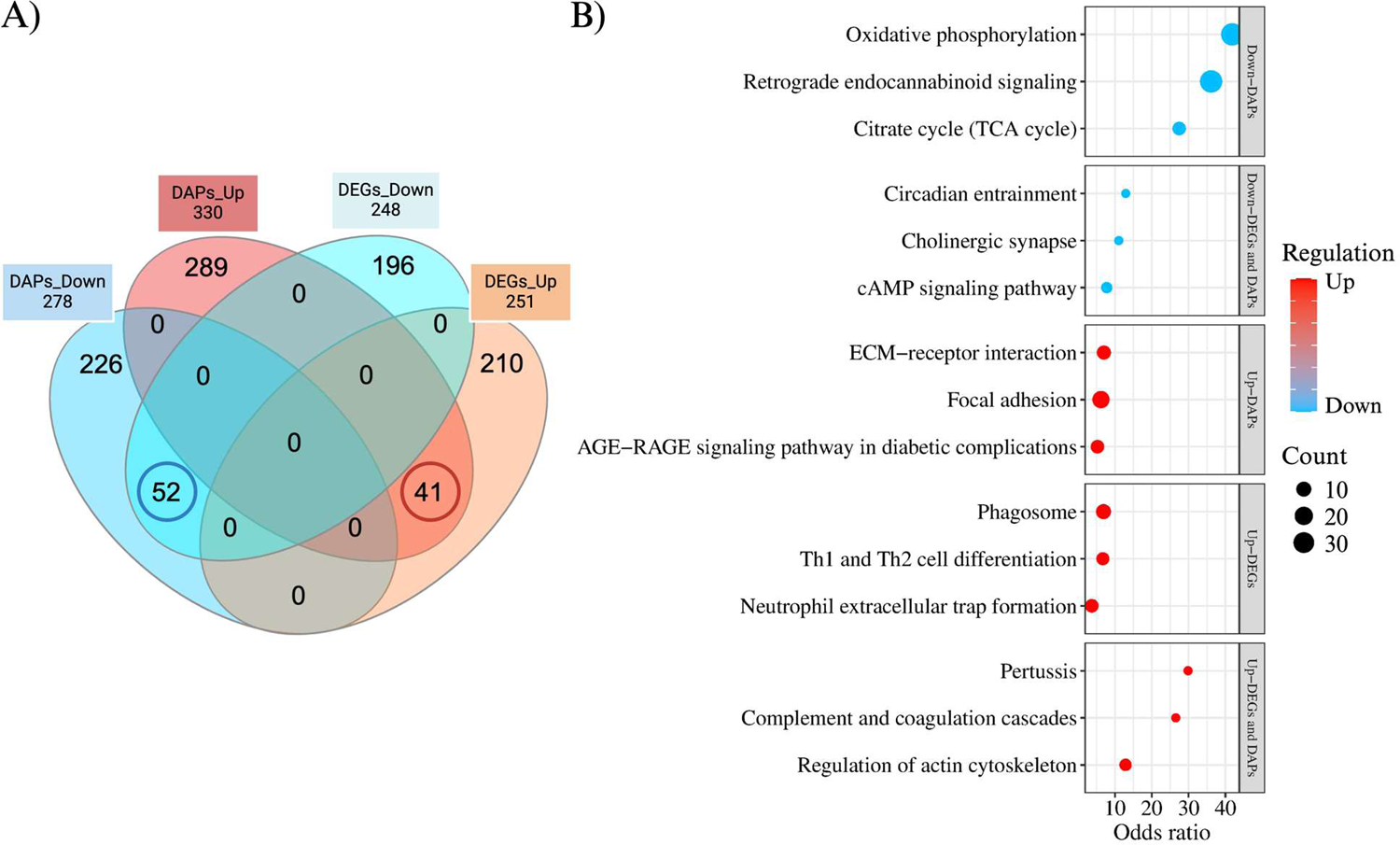
The comparison of robust DEGs and robust DAPs. A) Venn Diagram plot showing the common and unique changes between robust DEGs and DAPs; B) GO analysis displaying KEGG pathways for commonalities and differences between DEGs and DAPs. DEGs, Differentially Expressed Genes; DAPs, Differentially Abundant Proteins; GO, Gene Ontology.

**Table 2.** Characteristics of the selected transcriptome datasets included in this study.

| Source | Number. Cases/CTRs | Age (yrs.) AD/CTR | Post-mortem interval (hours) AD/CTR | Disease | Number of down/up-regulated genes | Ref. |
| --- | --- | --- | --- | --- | --- | --- |
| GSE118553 | 52/27 | 82.9 ± 8.7/<br>70.6 ± 15.9 | 39.9 ± 21.3/<br>37.1 ± 20.7 | AD | 268/396 | 29 |
| GSE48350 | 15/39 | 85.7 ± 6.3/<br>64.8 ± 9.5 | - | AD | 1009/663 | 30 |
| GSE33000 | 310/157 | 80.6 ± 9.0/<br>63.5 ± 9.9 | 13.7 ± 7.4/ 22.4 ± 5.8 | AD | 351/342 | 31 |

We then performed KEGG pathway enrichment analysis for these commonly altered transcripts and proteins. The complement and coagulation cascade, pertussis and regulation of actine cytoskeleton pathways were among the top up regulated pathways, while cAMP signaling, circadian entrainment and cholinergic synapse pathways were among the top down regulated pathways (**Figure 2B**). Results of KEGG pathway analysis revealed that most of the significantly enriched pathways for DAPs were not overlapped with pathways enriched for DEGs, specially for down regulated pathways. Notably, significantly downregulated proteins were enriched for oxidative phosphorylation and TCA cycle pathways (**Figure 1C**), although, these pathways did not show any changes at transcript level. These findings highlight the fact that AD brain mediates molecular changes at both transcript and protein level and analyses of discordance between transcript and protein levels can give rise to better understanding of molecular causes of AD as it was also shown previously ^32,33^.

### Cargo protein contents of large and small *EV*-enriched *subpopulations* showed significant overlap with the altered proteins in AD brains

To investigate whether dysregulated protein homeostasis in AD brain can be compensated with EVs, we analyzed the cargo protein content of different EV subpopulations derived from mesenchymal stem cells. Dependent on the isolation methods, which gave rise to different size distribution (see methods), EVs were classified into three different subpopulations, EV subpopulation isolated with high-speed centrifuge (HS-EVs), ultracentrifuge (UC-EVs) and ultracentrifugation on sucrose cushion (SU-EVs). We identified 2919 proteins commonly in all subpopulations (**Figure 3A**). Then, we analyzed the differences in abundance of each of these proteins across three EV subpopulations. The comparison of the protein abundances between HS versus UC group showed the highest (1103 high abundant, 401 low abundant, and 1415 proteins with no significant changes) and SU versus UC showed the lowest number (558 high abundant, 392 low abundant and 1969 no proteins with no significant changes) of proteins differentially abundant between them (**Figure 3A**). In addition, principal component analysis (PCA) showed the existing variation between protein abundances of each EV subpopulation (**Figure 3B**).

**Figure 3.**
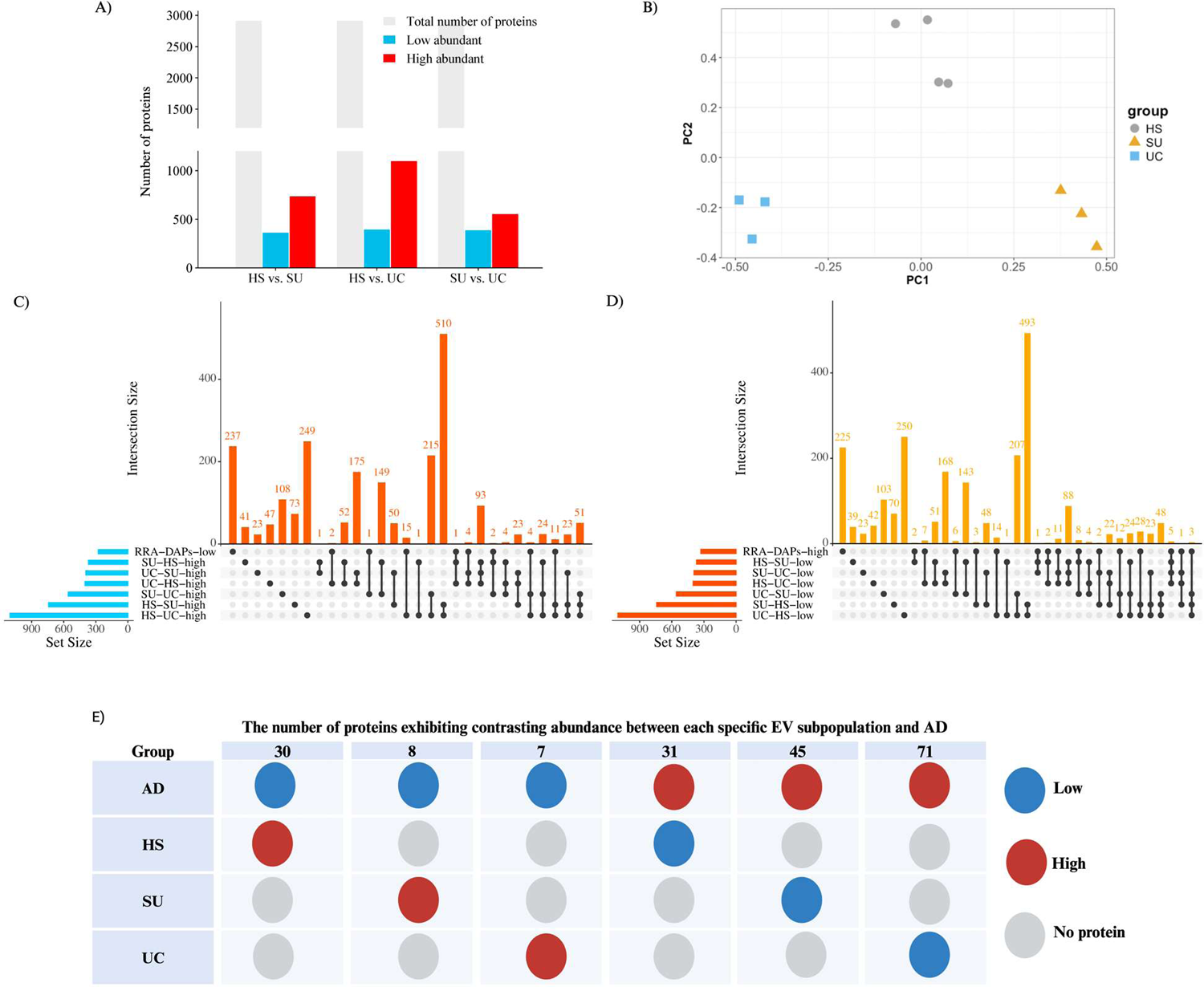
The binary comparison of differentially abundant proteins between EV subpopulations and their shared proteins with AD brains. A) The number of total, low and high abundant proteins in different EV subpopulations; B) The PCA analysis depicting the proteome variation between three different subpopulations of EVs; C) The comparison of DAPs with low abundance in AD with those DAPs with high abundance in EVs; D) The comparison of DAPs with high abundance in AD with those DAPs with low abundance in EVs; E) Number of proteins that showed contrasting abundance between EVs subpopulation and AD brains. DAPs, differentially abundant proteins; AD, Alzheimer’s disease; EVs, Extracellular vesicles; PCA, Principal component analysis.

Next, we investigated whether these differentially abundant EV cargo proteins from each comparison groups overlap with the altered proteins that we identified in AD brains. Our analyses revealed significant number of shared proteins (**Table 3, and Supplementary File 1**). Then, we identified which specific EV subpopulation that had the protein content exhibited the most significant opposite abundance in comparison to the protein level changes in AD brains (contrasting abundance; low in AD brain-high in EV subpopulation) (Figure 3C and 3D). For example, our analyses revealed higher abundance of 30 proteins in HS subpopulation that these proteins were characterized with significantly decreased abundances in AD brain in comparison to the healthy controls (**Figure 3E**).

**Table 3.**
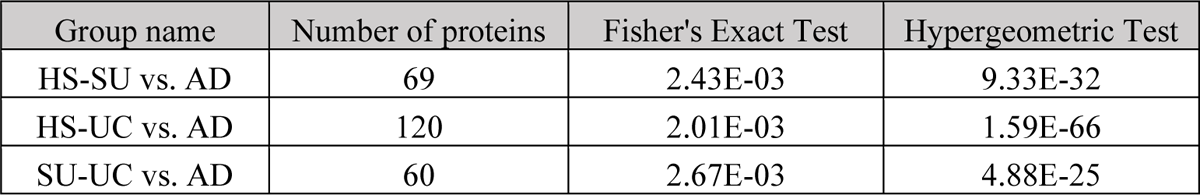
The Statistical Significance of observed protein commonalities between EV subpopulations and the AD brain was analyzed by Fisher’s exact test and Hypergeometric test.

### Pathway analysis revealed the HS-EVs as the potential most beneficial option for AD

Considering the idea of compensating dysregulated pathways associated with decreased proteins in AD brain as well as decreasing the risk of undesired effects of EVs, we further investigated the therapeutic potential of EV subpopulation through analyzing the biological function of DAPs specific to each EV subpopulation that showed contrasting abundance in AD brain.

To do this, list of DAPs obtained from Figure 3E were subjected to GO analysis including GO biological pathway, GO molecular function and GO cellular component analyses.

Results of functional enrichment analysis revealed unique and overlapping biological pathways for each EV subpopulations (**Figure 4**). Although, several biological pathways between different EV subpopulations, the number of proteins involved in those pathways were significantly different among them (**Figure 4, Supplementary file 2**). For instance, mitochondrial related pathways were identified in all three EV subpopulations, however, the number of proteins is different, where HS-EV group has the highest number of highly abundant mitochondrial proteins (**Figure 4A**) compared to SU (**Figure 4B**) and UC (**Figure 4C, Supplementary file 2)** subpopulations. As a result, the oxidative phosphorylation pathway showed the highest enrichment score in HS (258.98) compared to the SU (50.54) and UC (25.08) subpopulations **(Supplementary file 2)**. Another enriched pathway by high abundance proteins was TCA cycle that was enriched by DAPs in HS and SU EVs but not UC (**Figure 4, Supplementary file 2**). It should be highlighted here that in addition to the decreased mitochondrial function, TCA cycle is one of the top enriched pathways by low abundance proteins in AD brains (**Figure 2B).** Our findings suggests that these EV populations might be utilized to compensate decreased protein abundance for regulating TCA cycle in AD brain.

**Figure 4.**
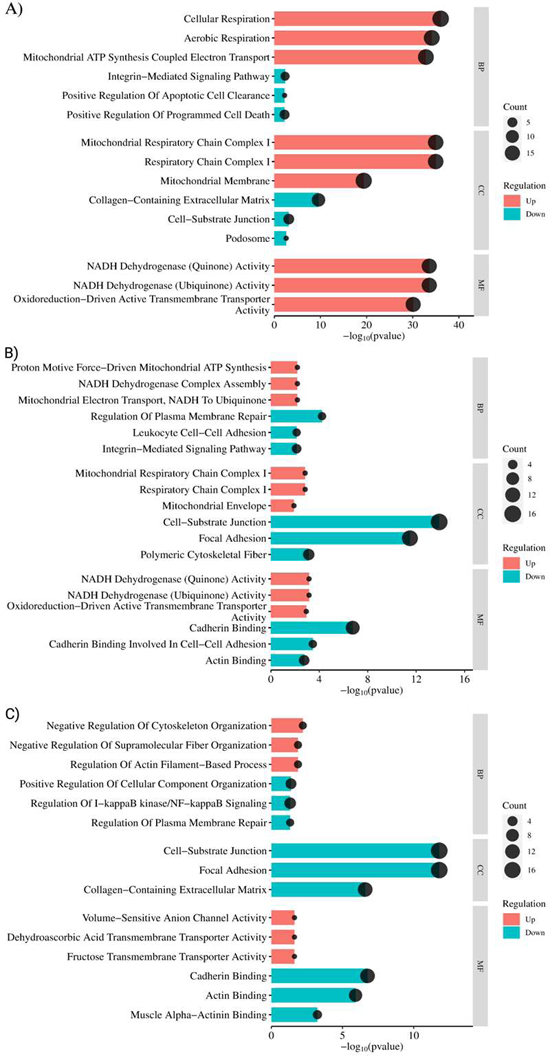
Gene ontology enrichment analysis of DAPs showed different directions of changes between EV subpopulations and AD brains. GO terms enriched for DAPs unique to; (A) HS-EVs, (B) SU-EVs, and (C) UC-EVs. GO, Gene Ontology; DAPs, differentially abundant proteins; EV, Extracellular vesicle; HS-EVs, high-speed centrifugation isolated extracellular vesicles; SU, Sucrose gradient ultracentrifugation isolated extracellular vesicles; UC, ultracentrifugation isolated extracellular vesicles.

These findings indicated that the heterogeneity of EVs in part can be attributed to differences in their cargo protein composition, and as such, they can be classified into distinct populations for potential application in clinical treatments. These observations also explain the inconsistency among published resulted regarding beneficial effects of different EV subpopulation.

Among top enriched pathways by low abundance proteins included, complement and coagulation cascades in HS group, but not other EV subpopulations (**Supplementary file 2**). Overall comparison of the enriched pathways between different EV subpopulations and AD brains revealed that HS-EV population has more related pathways to AD with opposite regulation, therefore may be a better therapeutic option for alleviating molecular changes in AD brains.

### Mitochondrial Energy Metabolism Proteins Central in HS-EVs Protein Network

To further investigate the potential therapeutic mechanism of HS-EVs to alleviate molecular changes in AD brains, we further expended our pathway analysis by including KEGG and Reactome tools that both yielded oxidative phosphorylation and complement and coagulation cascade as the top pathways enriched by high and low abundant proteins respectively (**Figure 5A and 5B**). Interestingly, out of 18 mitochondrial DAPs found in HS population, 16 of them were related to complex I of electron transport chain (ETC) (**Figure 5C**). The complex I specific dysfunction is implicated in different neurodegenerative disorders including AD and Parkinson’s disease (PD) and it’s been suggested as the main dysregulated complex resulting in mitochondrial dysfunction in AD ^34^.

**Figure 5.**
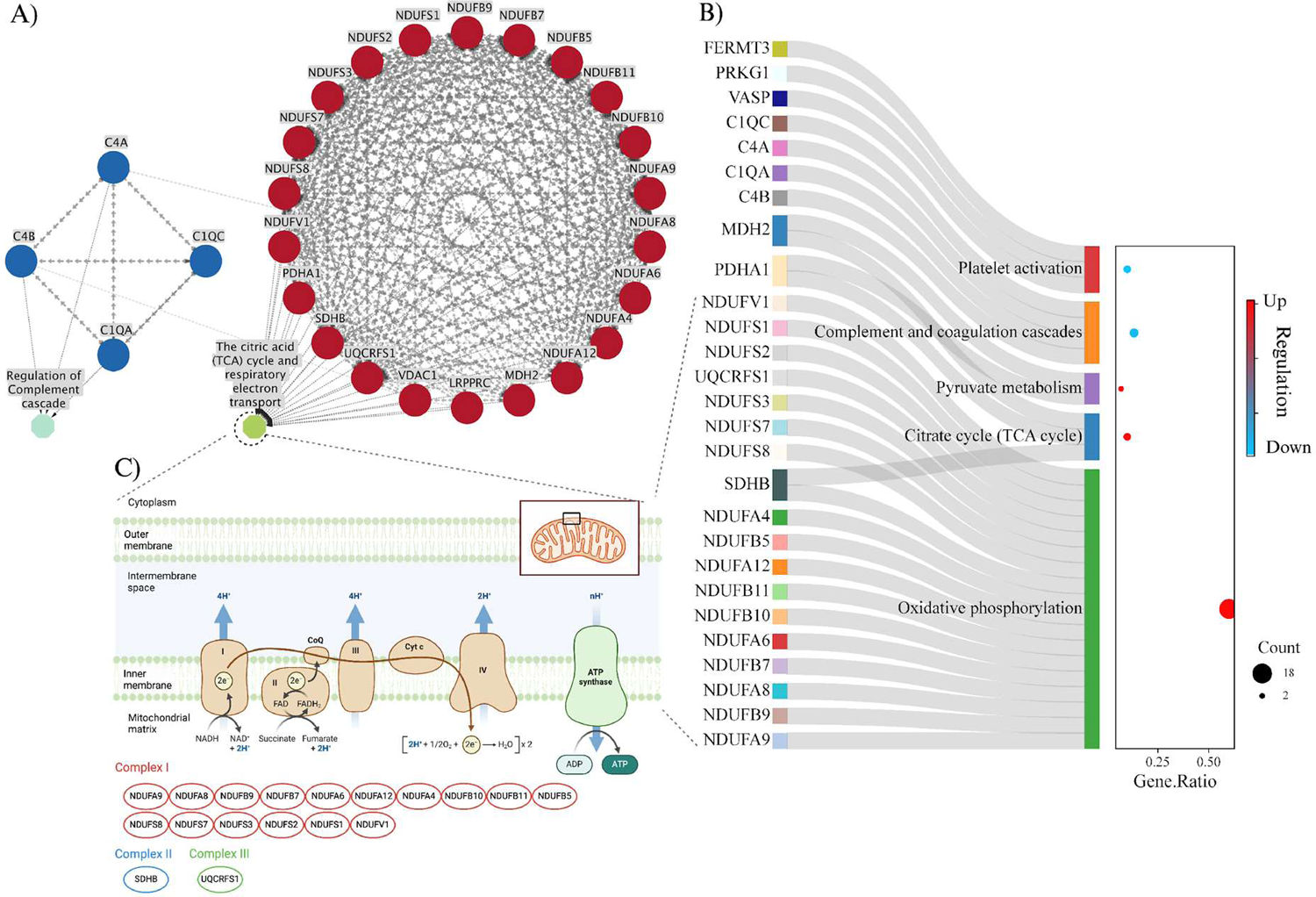
Analysis of proteins uniquely found in HS-EVs revealed mitochondrial pathways. A) Biological pathways obtained through Reactome database analysis; B) Biological pathways identified using KEGG database; C) The oxidative phosphorylation pathway (OXPHOS) as the primary pathway enriched by DAPs from the HS-EV group. Interestingly, 16 out of 18 enriched DAPs in OXPHOS are from complex I of ETC. HS-EVs, high-speed centrifugation isolated extracellular vesicles; AD, Alzheimer’s brain; ETC, electron transport chain.

### Hub Gene–miRNA-TF coregulatory network analysis revealed regulatory networks of miRNAs and TFs interacting with coding genes of key proteins within HS-EVs

Next, to examine the regulatory network of DAPs in HS population, CytoHubba plugin within Cytoscape was used, which yielded top 10 hub proteins. Intriguingly, 8 of these proteins were related to oxidative phosphorylation pathway including NDUFS8, NDUFB10, NDUFS3, NDUFS2, UQCRFS1, NDUFS1, NDUFV1 and SDHB (**Figure 6A**). There were also two other mitochondrial proteins, PDHA1 (Pyruvate Dehydrogenase E1 Subunit Alpha 1) and MDH2 (Malate Dehydrogenase 2) involved in tricarboxylic acid (TCA) cycle. PDHA1 is a subunit of the pyruvate dehydrogenase (PDH) complex that catalyzes the overall conversion of pyruvate to acetyl-CoA and CO_2_ and provides the primary link between glycolysis and the TCA cycle. The deficiency of PDHA1 exists in various diseases such as AD, epilepsy, Leigh’s syndrome, and diabetes-associated cognitive decline ^35^. The MDH2 has an oxidoreductase activity and its higher abundance in AD brains has been reported previously ^36^, while our results showed the lower abundance of this enzyme in AD brains. MDH2 and MDH1 is the mitochondrial isoform of Malate dehydrogenases (MDH) that catalyze the reversible oxidation of malate to oxaloacetate. MDH enzyme also has a key role in maintaining equilibrium of the NAD+/NADH ratio between the mitochondria and cytosol ^37^. Overall, these results further indicates that HS-EV subpopulation might have particular beneficial effect against dysregulated glucose metabolisms in AD brains.

**Figure 6.**
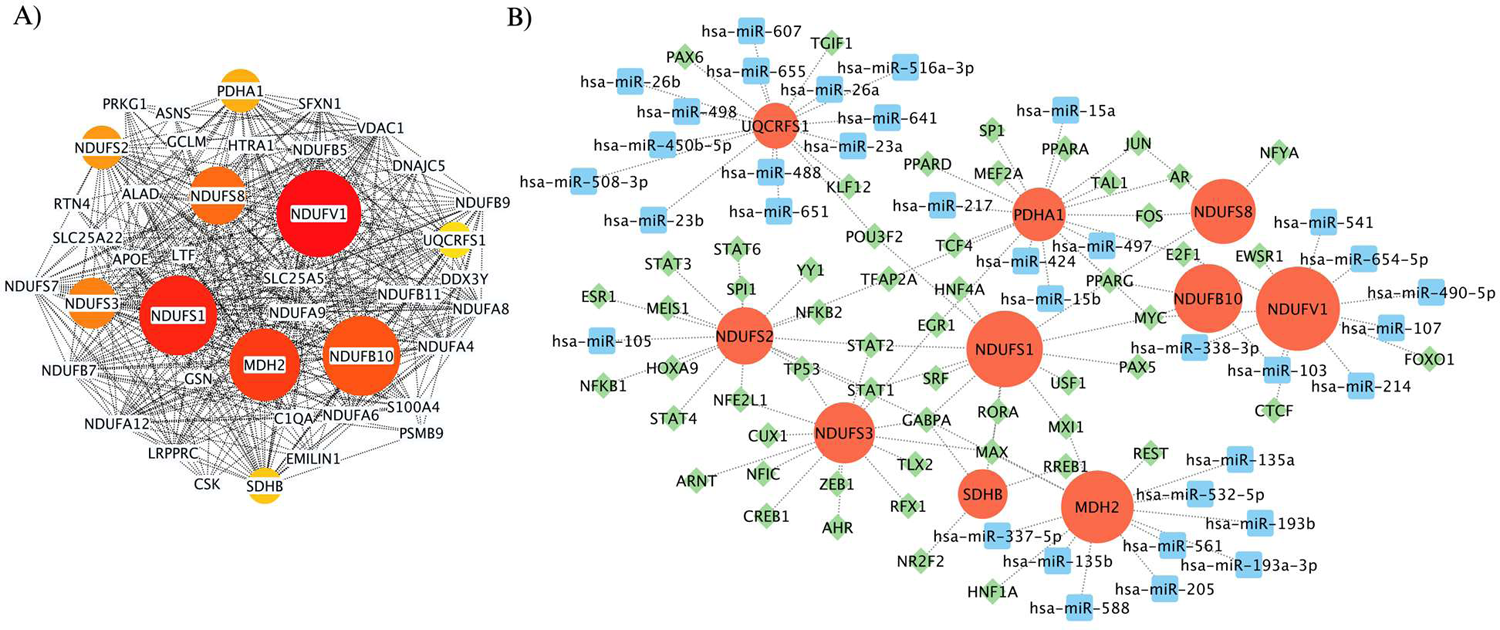
Regulatory network analysis of hub genes for DAPs uniquely found in HS-EVs. A) Top 10 hub genes within the interaction network: The size of the hub genes indicates their rank in the network based on the MCC algorithm. B) Coregulatory network of miRNA, TFs, and identified hub genes. DAPs; differentially abundant proteins; HS-EVs, high-speed centrifugation isolated extracellular vesicles; AD, Alzheimer’s brain; MCC, Maximal Clique Centrality algorithm.

Finally, to identify the upstream regulatory network of the hub genes, we identified miRNAs and Transcription Factors (TFs) regulating the expression of these hub genes (**Figure 6B**). Among the miRNAs that we identified, has-miR-103 and has-miR-484 was found to have highest connectivity score (see methods). For TFs, among the top candidates were GABPA, PPARG, and POU3F2 (**Table 4).** Next, to examine the relevance of these miRNAs and TFs to AD, we analyzed published papers by using a data mining approach using “easyPubMed” R package through PubMed database. The results of our data mining efforts revealed only a limited number of published studies investigating the relationship between these regulatory biological macromolecules (except for PPARG) and AD, highlighting an unknown mechanism underlying their association with AD (**Table 4, and Supplementary file 1**). While PPARG, a ligand-activated nuclear receptor that regulates lipid, glucose, and energy metabolism, has been found to exhibit altered levels in AD brains ^38^, the involvement of the other two transcription factors, POU3F2 and GABPA, in energy metabolism, cell proliferation, and differentiation, along with their roles in cognitive function and certain neurodegenerative diseases, lacks sufficient evidence regarding their association with AD^39,40^. Moreover, our analysis of the top identified miRNAs including has-miR-103 and has-miR-484, yielded similar outcomes, revealing a scarcity of data concerning the role of these miRNAs in the context of AD pathogenesis. Despite a few biomarkers discovery and bioinformatic studies suggesting a potential association of these miRNAs with neurodegenerative diseases, comprehensive understanding remains elusive ^41,42^.

**Table 4.** Number of published studies for top predicted regulators of hub genes.

|  | Target name | AD |
| --- | --- | --- |
| miRNA | has-miR-103 | 2 |
|  | has-miR-484 | 1 |
| TF | GABPA | 2 |
|  | PPARG | 610 |
|  | POU3F2 | 1 |

## Discussion

In this study, we performed a comparative-omics approach using the RRA method on proteome and transcriptome data of 788 human brains to obtain key altered genes and proteins and corresponding biological changes in AD. The RRA method utilizes fold change as the ranking criteria. In contrast to Venn diagram analysis, which identifies shared proteins, RRA identifies proteins that display significant fold changes across datasets, regardless of their presence in all datasets. In addition, we cross-compared the molecular changes regulated at transcript and protein levels in AD brains. Interestingly, while some common changes were observed at both transcript and protein levels, our results revealed more molecular changes at the protein level that were not captured at the gene expression level. These findings implicate the molecular deregulation beyond the gene expression level in AD, which may be mainly at protein level in both cellular processes in AD brain. Among the pathways that are altered at both transcript and protein levels, downregulation of cAMP signaling, and upregulation of complement and coagulation pathways were notable. The cAMP signaling pathway is integral to numerous biological processes. The cyclic AMP-protein kinase A-cAMP response element-binding protein (cAMP/PKA/CREB) pathway stands out as a prominent drug target in AD, as evidenced by a wealth of literature documenting the downregulation of cAMP signaling and CREB-mediated transcriptional cascades in AD^43^. This pathway is evolutionarily conserved across species. The cAMP response element-binding protein (CREB) serves as a widespread and intricately involved transcription factor that governs various aspects of neuronal function, including growth, differentiation, proliferation, synaptic plasticity, neurogenesis, neuronal maturation, spatial memory, long-term memory formation, and neuronal survival ^44^. Interestingly, the aberrant energy metabolism was one of the main findings identified at the protein but not at the gene expression level. This data indicates translational dysregulation of mitochondrial energy metabolisms. Deregulated metabolic pathways such as OXPHOS and TCA cycle have been widely investigated in AD pathogenesis and have been shown to contribute to the progression of the AD ^45^. The differences between gene expression and protein translation changes in AD brains have been previously reported ^32,33^. In line with our findings, it was shown that protein modules and disease-associated expression changes at protein translation levels do not directly mirror changes at the mRNA level in AD. Changes in the mitochondrial protein alterations was described as one of the earliest events in AD, and these changes were found to be more evident at the protein level^32^. In conjunction with a few previously published reports, our findings indicate a stronger association between protein level alterations and dysregulated cellular processes in the AD brain.

In the next step, to investigate the potential of different subpopulations of EVs as therapeutic agents against dysregulated pathways in AD brain, we carried out a comparative analysis of proteome data by comparing the proteins identified in different MSCs-derived EV subpopulations to proteome data from AD brains. There is mounting experimental evidence indicating size-dependent beneficial effects of different EV subpopulation. While in some reports beneficial effects of small EVs have been shown^11,12^, others indicated the superior beneficial effects of large EVs for alleviating molecular phenotypes of AD ^13^. These observations were attributed to the size and size distribution of EV subpopulation, ultimately effecting their cargo contents and their therapeutic properties ^14,15^. The size and size distribution of EVs is mainly affected by the isolation methods. Therefore, in this study we have analyzed and compared the molecular properties of proteome data from three different EV subpopulations isolated through different methods.

Our study highlighted the EV differences in terms of their application as a potential treatment for AD, where EVs isolated through high-speed centrifugation with higher size distribution showed to be the best option for AD. HS-EVs turned out to have more proteins involved in energy metabolism like proteins involved in OXPHOS and TCA cycle, while fewer proteins involved in inflammatory responses like regulation of complement and cascade pathways. Interestingly, results of hub gene analysis using MCC on DAPs in HS-EVs also returned the mitochondrial proteins including NDUFV1, NDUFS1, MDH2, NDUFB10, NDUFS8, NDUFS3, NDUFS2, PDHA1, SDHB and UQCRFS1 as the key genes. We used MCC method that focuses on maximal cliques, which are subsets of genes where each gene is directly connected to every other gene in the subset. By analyzing these maximal cliques, MCC identifies genes that are densely connected within the network, indicating their potential importance in regulating biological processes or pathways. This approach helps uncover central players or hub genes that may have significant influence or control over molecular interactions and cellular functions and suggested to be the best method for hub gene prediction ^19^.

Aberrant energy metabolism is one of the key pathological events involved in progression of many neurodegenerative diseases including AD, Parkinson’s disease and Huntington’s disease^46^. Impaired energy metabolism occurred in AD, even before clinical symptoms arise and is driving cognitive dysfunction ^47^. Glucose is the main source of energy for brain under physiological conditions, and dysregulation of energy-related metabolisms such as TCA cycle and oxidative phosphorylation in some brain regions including frontal cortex is correlated with cognitive decline^46^. Mitochondria is the supplier of energy in the brain which it’s impairment and consequently energy deficient leads to neuronal death and neurodegeneration ^48^. It’s been reported that inhibition of OXPHOS/ETC activity activates microglia leading to the elevation of pro-inflammatory cytokine production and neuroinflammation ^49^. Neuroinflammation is one of the pathological features of AD, rapidly consuming glucose and depleting energy to the neurons ^46^. Mounting evidence on the key role of dysfunctional mitochondria in disease initiation and progression has led to the idea of mitochondria transplantation from healthy donor to affected recipients as a potential therapeutic intervention for many diseases ^50^. This approach opened a new window in the treatment of metabolic diseases like AD, and a growing number of studies have reported promising results of mitochondrial transplantation for neuronal regeneration ^50,51^. However, transferring the whole mitochondria is still challenging because of critical concerns such as isolation of undamaged and coupled mitochondria since damaged mitochondria may exacerbate the disease, denser mitochondria may mean detrimental ROS amplification and more cytochrome C produced when cells are exposed to a harsh environment leading to insufficient cell clearance and cell apoptosis, and transplant rejection ^51,52^. Therefore, transferring only part of the mitochondria or proteins may be a better option that need further investigations. Recently, mitochondrial vesicles referred as mitovesicles, isolated using ultracentrifugation (100,000g), showed to secret from mitochondria and contain a specific subset of mitochondrial constituents that showed changes between healthy and disease condition^53^, however their functions and potential applications remained to be elucidated. Interestingly, our analysis showed that most of the high abundant mitochondrial proteins in HS group were related to complex I of ETC, which is a main affected part of ETC in neurodegenerative diseases particularly AD ^54^. Approximately 99% of mitochondrial proteins are encoded by nuclear genes, mainly synthesized as precursor proteins on cytosolic ribosomes and imported into mitochondria through different ways and become mature^55^. Mitochondria need energy to convert the immature proteins to mature proteins, which lack of enough energy in disease condition may affect this process, in addition, Aβ peptides can permeabilize the mitochondrial membrane and disrupt the maturation of mitochondrial proteins ^56^. Interestingly, the comparative transcriptome analysis of AD brain and healthy controls did not show the down regulation of oxidative phosphorylation pathway, indicating the disturbance is mainly in the translational level^18^. Besides, ETC proteins, our analysis showed the higher abundance of TCA cycle proteins including PDHA1, MDH2 and SDHB in HS-EVs than other EV subpopulations. Decreased expression of PDHA1 has been reported in neurodegenerative diseases such as AD and PD, and conditional knockout of PDHA1 led to cognitive function impairment in mice ^35,57^. PDHA1 serves as a pivotal component of the pyruvate dehydrogenase complex (PDC), crucial for catalyzing the oxidative decarboxylation of pyruvate into acetyl-CoA, which connect the cytoplasmic glycolytic pathway with the mitochondrial TCA and oxidative phosphorylation ^58^. The lack of PDC results in inadequate energy provision to the brain since neuronal ATP generation primarily occurs in the mitochondria through the oxidative phosphorylation of glucose via the TCA ^35^. On the other side, PDC deficiency causes an elevation in lactate levels which can induce oxidative stress and apoptosis in cortical neurons ^35^. An age-related increase in lactate levels has been reported in APP/PS1 mice that correlated with impaired memory performance ^59^. MDH2 is another TCA related proteins that showed to has low abundant in AD brains but was high abundant in HS-EVs. MDH is the final enzyme in the mitochondrial TCA cycle that catalyze the inter-conversion of L-malate and oxaloacetate using nicotinamide adenine dinucleotide (NAD) as a cofactor to generate reducing equivalents ^36^. Deficiency in MDH2 results in psychomotor delay, muscular hypotonia, recurrent seizures and pediatric epileptic encephalopathy. These symptoms stem from decreased energy production in the brain and skeletal muscles, both of which require substantial energy resources ^37,60^. Interestingly, increased mRNA level of MDH2 in AD cases compared to healthy controls has been reported ^36,61^, indicating that the disruption in the AD may be at translational level leading to low abundance of the MDH2 at protein level. SDHB is another protein involved in TCA that showed to be low abundant in AD brains and high abundant in HS-EVs. SDHB, which encodes the iron-sulfur subunit B of succinate dehydrogenase (SDH or complex II), facilitates the transfer of electrons from flavin adenine dinucleotide (FADH) to coenzyme Q during succinate oxidation ^62^. Down regulation of SDHB in the mRNA level has been reported previously and its low amount at protein level has been linked to cognitive decline following estrogen deficiency ^63^. The presence of mitochondrial proteins in stem cell derived EVs and their beneficial effects in restoring mitochondrial function and decreasing the pathological events in different diseases including AD have been previously reported ^64–67^.

The comparison of HS-EVs vs other EV subpopulations not only revealed higher abundance of beneficial proteins for AD, it also returned lower abundance of proteins that showed to have increased level in AD and contribute to its pathogenesis. Of most important proteins are proteins involved in inflammatory responses such as complement and coagulation cascade pathway including C1qA, C1qB, C4B and C4A, that has been widely investigated in the pathogenesis of AD^18^. The C1q is a subcomponent of the classical pathway of the complement system, it is a complex protein made up of three different polypeptide chains: C1qA, C1qB, and C1qC. Previous reports indicated that in the presence of the fibrillar plaque pathology, C1q cause a detrimental effect on neuronal integrity, possibly via triggering the activation of classical complement cascade, while lack of these proteins led to less neuropathology in AD mice model ^68^. Increased expression of C1A and C1B also have been reported in AD patients and showed to be associated with AD-related neuropathology ^69^. Intriguingly, level of C1A and C1B showed to be different in control groups based on the Apolipoprotein E genotype with significantly lower level in ε2/ε3 subjects compared to ε3/ε3 and ε3/ε4 subjects, led to growing hypothesis of protective effects of APOE ε2 for AD ^69,70^.

Next, we have performed a comprehensive miRNA and TF analysis on the yielded hub genes and obtained the key miRNAs and TFs that may regulate the expression of hub genes. Data mining results revealed that, in sipe of the strong connection between identified miRNAs and TFs with hub genes, their role in AD is not well understood and required further investigations. The has-miR-103 has been reported as a potential plasma biomarker for AD detection, and it has been showed to interact with NDUFV1 gene and possibly leads to decreased expression of this gene^41,71^. On the role of has-miR-484, although previous bioinformatic analysis showed the enrichment of this miRNA in the molecular changes associated with AD, the level of this miRNA was identified as the most stable reference genes in the plasma samples of 38 healthy controls, 26 MCI, 56 AD, and 27 frontotemporal dementia (FTD) patients and was used as the reference for the Quantitative real-time PCR (RT-qPCR) data normalization ^42,72^. Among identified TF, GABPA (also known as nuclear respiratory factor 2 (NRF 2), a member of the E-twenty-six (ETS) family of DNA-binding factors, plays a pivotal role in regulating the expression of many genes with roles in cell cycle control, apoptosis, differentiation, hormonal regulation, mitochondrial functions, and other essential cellular processes ^40,73^. GABPA also regulates nuclear genes responsible for encoding mitochondrial proteins and showed to be down regulated in brain tissues that may led to mitochondrial dysfunction ^74^. Moreover, many genes involved in AD have been reported to have a GABPA binding site ^75^, however its role in AD pathogenesis is not well understood. POU3F2 is another key TF interacting with hub genes, which has a role in regulation of proliferation and differentiation of neural progenitor cells (NPCs). Deregulation of this gene showed to decrease NPC differentiation toward neurons, increase NPCs proliferation and disrupt the excitatory synaptic transmission of NPC-derived neurons ^39^. Down regulation of POU3F2 has been reported in postmortem brain tissue from patients with Schizophrenia and bipolar disorder ^76^. POU3F2, also showed to be involved in cognitive function and adult hippocampal neurogenesis, and mice lacking its expression exhibited impairment in cognitive function, yet its role in AD is not clear ^77^. The last key TF that was identified in our analysis is PPARG, which has been widely studies in brain function and neurodegenerative diseases. PPARG (PPARγ), a member of the nuclear receptor family Peroxisome proliferator-activated receptors (PPARs), serves as a transcription factor governing the expression of genes pivotal in adipogenesis, lipid metabolism, inflammation, and the maintenance of metabolic homeostasis ^38^. Notably, PPARγ exerts a suppressive effect on BACE1 expression via its responsive element present in the promoter region, thereby inhibiting Aβ production ^78^. Studies have demonstrated that activation of PPARγ can effectively attenuate the expression of inflammatory cytokines, iNOS, and NO production in microglial cells, while also inhibiting COX2 activity, thus indicating antagonistic effects against the transcription factor NFκB ^79^. Furthermore, in a rat model of cortical Aβ injection, the administration of PPARγ agonists such as ciglitazone, ibuprofen, and pioglitazone significantly suppressed Aβ-induced microglial cytokine production, highlighting the potential of PPARγ as a therapeutic target for Alzheimer’s disease ^38,80^. In our gene regulatory network analysis, we uncovered both known and novel genetic components of biological pathways linked to AD, shedding light on the potential mechanisms underlying the therapeutic effects of various EV subpopulations. Beyond the utilization of EVs, the genes, proteins, miRNAs, and TFs identified in our study offer promising targets for therapeutic intervention aimed at reducing the risk, delaying the onset, or mitigating the progression of AD.

## Conclusions

In conclusion, our research provides significant insights into the molecular alterations occurring in AD at both the gene and protein levels, while also elucidating the potential therapeutic properties of distinct EVs in combating this debilitating condition. Through comprehensive transcriptomic and proteomic analyses, we identified common transcriptional and proteomic signatures in the frontal cortex of AD-affected individuals, highlighting protein-level changes not apparent at the gene level. This illuminated key pathways affected in AD progression, particularly revealing mitochondrial dysfunction and impaired energy metabolism pathways such as dysregulated oxidative phosphorylation and the TCA cycle, which could be targeted by EV cargo.

Furthermore, our study pinpointed several promising therapeutic targets for AD, including genes, proteins, miRNAs, and TFs. Restoring mitochondrial function and energy metabolism emerges as a potential neuroprotective strategy, while modulating inflammatory responses and targeting specific hub genes, proteins, miRNAs, and TFs offer prospects for alleviating AD symptoms. However, it is essential to emphasize that the therapeutic targets identified require experimental validation to ensure their clinical efficacy and safety. Therefore, further investigation utilizing animal models and large-scale clinical cohorts is imperative to validate and build upon our findings.

## Acknowledgments

We would like to acknowledge the generous funding provided by the NIA/NIH (1K01AG060040) and The Jeffress Trust Foundation to AK.

## Author contributions

Conceptualization; Abyadeh M., and Kaya A. Data Curation and Formal Analysis; Abyadeh M. Writing – Original Draft; Abyadeh M., and Kaya A. All authors have approved the final article.

## Supporting Information

**Supplementary File 1:** Supplementary File 1 includes proteomics and transcriptomics datasets for AD brain samples and EVs, as well as DAPs and DEGs resulted from the analyses of each dataset, along with normalized counts and their p values. It also includes a cross-comparison of DEGs and DAPs.

**Supplementary File 2:** GO enrichment terms along with statistical significance for DEGs, DAPs, proteins, and transcripts commonly shared by both datasets identified in AD brain samples and EV subpopulations.

